# CG14906 (*mettl4*) mediates m6A methylation of U2 snRNA in *Drosophila*

**DOI:** 10.1101/2020.01.09.890764

**Authors:** Lei Gu, Longfei Wang, Hao Chen, Jiaxu Hong, Zhangfei Shen, Abhinav Dhall, Taotao Lao, Chaozhong Liu, Zheng Wang, Yifan Xu, Hong-Wen Tang, Damayanti Chakraborty, Jiekai Chen, Zhihua Liu, Dragana Rogulja, Norbert Perrimon, Hao Wu, Yang Shi

## Abstract

Recent studies reported that METTL4 regulates DNA 6mA in vivo and therefore is a candidate DNA m6A methyltransfease. However, the enzymatic activity of METTL4 in vitro has not been demonstrated in part due to the difficulties of obtaining well-folded proteins. Here we show that mettl4 is a major methyltransfase responsible for m6A methylation of U2 snRNA both in vitro and in vivo in fly, and identify adenosine at 29th position as the site of m6A methylation. This study answered a long-standing question regarding the enzymatic activity of METTL4, and thus paved the way for further investigating the functions of METTL4 in different biological settings.

## Introduction

While the eukaryotic candidate m6A methyltransferases belong to multiple distinct methylase lineages, the most widespread group belongs to the MT-A70 family exemplified by the yeast mRNA adenine methylase complex Ime4/Kar4. At the structural level, all of these enzymes are characterized by a 7-β-strand methyltransferase domain at their C-terminus, fused to a predicted alpha-helical domain at their N-terminus and require S-adenosyl-L-methionine (SAM) as a methyl donor. The catalytic motif, [DSH]PP[YFW], shown to be critical for METTL3/METTL4-mediated mRNA m6A methylation[1]. The high degree of amino acid sequence conservation among the predicted N6-Methyladenosine methyltransferases motivates further explorations into their potential functional conservation. METTL4 is a member of the MT-A70-like protein family, which is conserved during evolution (**Fig.1a**) [2]. Previous studies suggested that METTL4 regulates DNA 6mA *in vivo* and therefore is a candidate DNA m6A methyltransfease [3–5]. However, the enzymatic activity of METTL4 *in vitro* has not been demonstrated.

**Fig.1.**
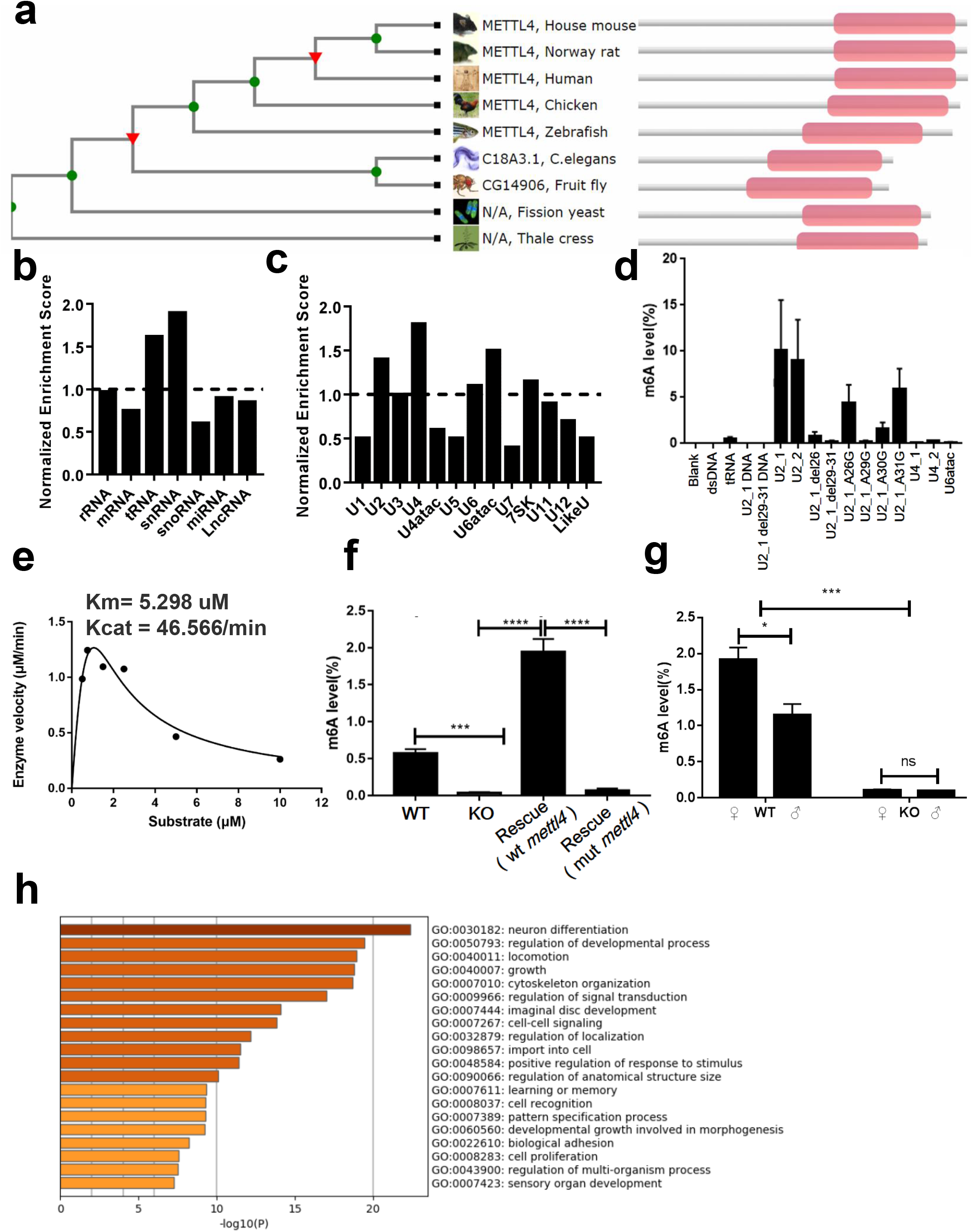
CG14906 (*mettl4*) methylates U2 snRNA in *Drosophila melanogaster*. **a.** Cladogram of mettl4 in model organisms based on their sequence similarity, the pink rectangle indicates the MT-A70 domain. **b.** Enrichment analysis of eCLIP-seq data for different RNA types. tRNA and snRNA are enriched among all RNA types in general and snRNA is the top enriched RNA molecules targeted by mettl4 *in vivo*. **c.** Enrichment analysis of eCLIP-seq data for subgroups of snRNA. U2, U4, and U6atac are the top enriched subgroups. **d.** *In vitro* enzymatic activity is measured by LS-MS/MS using substrates including U2, U4, U6atac, U2 with different point mutations and deletions, tRNA and DNA with U2 sequences. Results show that U2 is the best substrate for fly mettl4 and that adenosine at position 29 in U2 is methylated by METTL4. **e.** Michaelis-Menten kinetics of recombinant *mettl4* was determined using U2 probes as substrate by LC-MS/MS analysis. **f.** U2 m6A analysis in WT, KO and rescued (wt: wild type *mettl4*; mut: catalytic dead mutant *mettl4*, DPPW→NPPW) cells by LC-MS/MS. **g.** *In vivo* U2 m6A analysis by LC-MS/MS of WT and KO flies. Error bars indicate mean ± s.d. (n=3). **h.** Genes with differential alternative splicing were used for the GO analysis. The top 20 enriched biological processes are shown in the bar plot. Statistical significance is determined as: ns = p > 0.05; * = p < 0.05; ** = p < 0.01; *** =, p < 0.001; **** = p < 0.0001.

### Identification of potential substrates by eCLIP-seq

To identify the substrate(s) for METTL4, we purified his-tagged, wildtype as well as a catalytic mutant (DPPW mutated to NPPW) (Fig. S1) of *Drosophila melanogaster mettl4* from *E. Coli* strain BL21 (DE3). In order to unbiasedly identify potential substrates of *mettl4*, we performed *in vitro* enzymatic assays using various substrates including both DNA and RNA with and without secondary structures. We used deuterated S-Adenosyl methionine (SAM-d3) in the in vitro enzymatic assays in order to identify m6A mediated by mettl4. Although we detected a weak enzymatic activity on DNA substrates composed of previously published sequence motifs, *mettl4* appears to prefer RNA substrates with secondary structures (**Fig.S2**). We next performed eCLIP-seq, which was originally developed to map binding sites of RNA Binding Proteins on their target RNAs [6], to identify the RNA type that is targeted by *mettl4 in vivo*. Since there are no commercial antibodies available for fly *mettl4*, we generated a *Drosophila* KC cell line with a FLAG-tagged *mettl4* for the eCLIP-seq experiment [7]. In total, we generated two biological replicates for immunoprecipitation (IP) samples, and their respective input samples, together with one IP-control and Input-control sample for the quality control and enrichment analysis [8]. The two replicates showed a strong correlation with a Spearman correlation coefficient of 0.97, indicating great consistency between the replicates (**Fig.S3**). Thus, we merged the two replicates to increase the sequencing depth and power for downstream analyses, which showed that *mettl4* captured RNA molecules, mostly tRNA and snRNA, including U2, U4 and U6atac (**Fig.1b, c**).

### Mettl4 catalyze U2 m6A *in vitro*

We next investigated whether the RNAs identified by the eCLIP experiments are indeed substrates of METTL4 by carrying out in vitro enzymatic assays. We synthesized oligonucleotides containing tRNA and snRNA sequences and various controls including DNAs with the same sequences. The *in vitro* enzymatic activity of *mettl4* on each candidate substrate and control sequences was measured by LC-MS/MS. These in vitro experiments led to the identification of U2 as the best substrate among all the snRNA subtypes (**Fig.1d**). Next, we wished to identify the adenosine residues in U2 that are methylated by METTL4. Previous studies documented that the adenosine at the 30th position of U2 is frequently methylated in vertebrate U2 snRNA [9], with a sequence motif of AA-G as opposed to _28_AAAG_31_ in fly. To identify which adenosine within the motif is essential for the enzyme activity in fly, we generated point mutations and deletions of adenosine within and close to this motif and measured the enzymatic activity of *mettl4* on these substrates. We found that when the 29^th^ position adenosine is mutated or deleted, no m6A methylation was detected by LC-ms/ms, whereas other point mutations or deletions (26^th^ and 31^st^ positions) did not affect substrate methylation or only decreased methylation partially (i.e., the 30^th^ position). These results indicate that adenosine at position 29 is the adenosine in U2 that is methylated by *mettl4* in fly (**Fig.1d**). In order to better characterize the enzymology of *mettl4*, we next investigated the kinetics of *mettl4* and determined that *mettl4* was able to methylate U2 with a Michaelis-Menten constant (K_m_) of 5.298 μM and a catalytic rate constant (k_cat_) of 46.566 min^−1^ (**Fig.1e**). In addition, the enzyme is inhibited by the substrate at higher concentrations (**Fig.1e**).

### Mettl4 catalyze U2 m6A *in vivo*

Next, we investigated whether mettl4 catalyzes U2 m6A *in vivo*. To accomplish this goal, we generated mettl4 KO and rescue cell lines (rescued by either wildtype or catalytic mutant of mettl4) (**Fig.S4** and **S5**). Indeed, the U2 m6A level is decreased dramatically in the *mettl4* KO cells and restored in the wt *mettl4* rescued cells, but not in the catalytic mutant *mettl4* rescued cells (**Fig.1f**, **Fig.S7 and Fig.S8a**). Furthermore, the same reduction of U2 m6A level was also seen in KO flies (**Fig.1g**, **Fig.S6, Fig.S7 and Fig.S8b**). The low DNA 6mA levels between WT and KO fly cells for both nuclear and mitochondrial DNA showed no significant differences (**Fig.S9**). These findings suggest that it is *mettl4* that mediates U2 methylation in vivo. Interestingly, the U2 m6A level in wildtype female flies is significantly higher than in males, suggesting that *mettl4* might play sex-specific roles (**Fig.1g** **and Fig.S8b**), which will be interesting to investigate in the future. Given U2 snRNA is involved in pre-mRNA splicing[10], we performed RNA-seq using both wild type and knockout *Drosophila* KC cell lines to determine if RNA splicing is affected as a result of mettl4 loss. In total, we identified 2,366 transcripts with differential alternative splicing events, which cover 1,771 genes. Gene Ontology Enrichment analysis suggests that *mettl4* affects a broad set of biological processes, including differentiation, development, growth, and response to stimulus (**Fig.1h**).

## Discussion

Since U2 is an essential component of the major spliceosomal complex, which plays an important role in pre-RNA splicing, loss of mettl4 might have broad impacts through altered RNA splicing. However, whether the altered RNA splicing events are regulated by *mettl4* through methylation of U2 snRNA or other yet-to-be-identified substrates, or whether *mettl4* regulates splicing in an enzymatic activity-independent manner, remain to be determined in the future.

In summary, we demonstrated that mettl4 catalyzes U2 m6A in fly both *in vitro* and *in vivo* and identified adenosine 29 in U2 snRNA as the site of methylation by mettl4. Furthermore, whole transcriptome profiling revealed that loss of *mettl4* broadly impacts various biological pathways. Our work answered a long-standing question regarding the enzymatic activity of *mettl4*, and thus paved the way for further investigating the functions of *mettl4* in different biological settings.

## Acknowledgements

We are grateful to all members of the Shi lab for general support and Drosophila RNAi Screening Center (DRSC) for excellent technical support. This work was supported by BCH funds and an epigenetic seed grant from Harvard medical school (601139_2018_Shi_Epigenetics). L.G is supported by NIH T32 award (4T32AG000222-25 and 2T32AG000222-26). Y.S. is an American Cancer Society Research Professor. D.R. is a New York Stem Cell Foundation-Robertson Investigator. This work was also supported by The New York Stem Cell Foundation.

## AUTHOR CONTRIBUTIONS

L.G., H.C. and Y.S. conceived, designed and coordinated the project. L.G. performed data analysis. L.G. and L.W., performed most of the *in vitro* experiments. J.H. generated *mettl4* KO cell line and performed *in vivo* analysis. A.D. performed kinetics analysis. Z.S., Z. W., and Y. X. helped the *in vitro* and *in vivo* experiments under the supervision of L.G., L.W., D. C., H.C., T. L., Z. L. and H.T. J.C. generated *mettl4* KO fly. N.P., D. R., H.W., and Y.S. supervised the project in general. L.G. and Y.S. wrote the manuscript with support from all authors.

## Figure Legends

**Figure S1 SDS-PAGE of recombinant proteins encoded by CG14906 (*mettl4*)**. His-tagged recombinant *Drosophila* melanogaster METTL4 proteins (wildtype and a catalytic mutant with a point mutation in the DPPW motif: DPPW to NPPW) were purified from the *E.coli* strain BL21 (DE3).

**Figure S2 *In vitro* enzymatic activity of Drosophila METTL4 on DNA and RNA substrates with different sequences.** Both DNA and RNA substrates, upon incubation with the recombinant METTL4, were measured for the m6A level by LC-MS/MS. A low level of m6A was observed in the DNA substrates and a relatively high level of m6A was observed in structured RNA substrates. Error bars indicate mean ± s.d. (n=3).

**Figure S3 Correlation between biological replicates of the eCLIP-seq samples.** Reads aligned to fly reference genome (dm6) were normalized in reads per million and the reads density for each 1-kb window was used for the Spearman correlation coefficient calculation.

**Figure S4 Generation of a knockout Kc cell line.** *mettl4* in fly Kc cell line was knocked out by **CRISPR-Cas9**. Small indels produced by a single guide RNA caused frame shift mutations and the editing efficiency was over 90% based on the ICE CRISPR Analysis Tool.

**Figure S5 Rescue of m6A level of U2 snRNA by overexpressing *mettl4* in the *mettl4* KO cells.** The Pentry-PAWF Gateway system was used to over-express *mettl4* in the fly Kc KO cells. The expression level of *mettl4* was measured by qPCR and normalized to Kc pAWF.

**Figure S6 Generation of *mettl4* knockout fly.** *mettl4* knockout flies were generated using the CRISPR/Cas9 system and verified by PCR.

**Figure S7 MS spectra of U2 in *mettl4* WT, KO and the rescued cells.** A, m6A and m6Am levels were measured by LC-MS/MS and compared among WT, KO and rescued cells. Standard peaks indicate the retention time for each modification.

**Figure S8 LC-MS/MS results for other independent KO cell lines and flies. a.** m6A levels for the other two independent KO cell lines and **b.** one KO fly line were measured by LC-MS/MS. Error bars indicate mean ± s.d. (n=3). Statistical significance is determined as: ns = p > 0.05; * = p < 0.05; ** = p < 0.01; *** = p < 0.001; **** = p < 0.0001.

**Figure S9 LC-MS/MS results for m6A on nuclear and mitochondrial DNA from WT and KO fly cells.** m6A levels for both nuclear and mitochondrial DNA from WT and KO fly cells were measured by LC-MS/MS. Error bars indicate mean ± s.d. (n=3). Statistical significance is determined as: ns = p > 0.05.

