## Supplementary material for "CG14906 (*mettl4*) mediates m6A methylation of U2 snRNA in *Drosophila*": Suppl fig

### Figure S1 SDS-PAGE of purified protein encoded by CG14906 (*mettl4*)

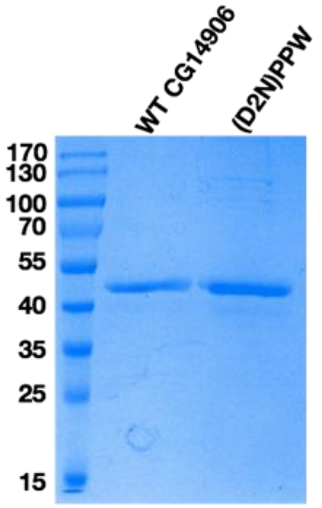

### Figure S2 *in vitro* enzymatic activity of *Drosophila mettl4* on DNA and RNA substrates with different sequences

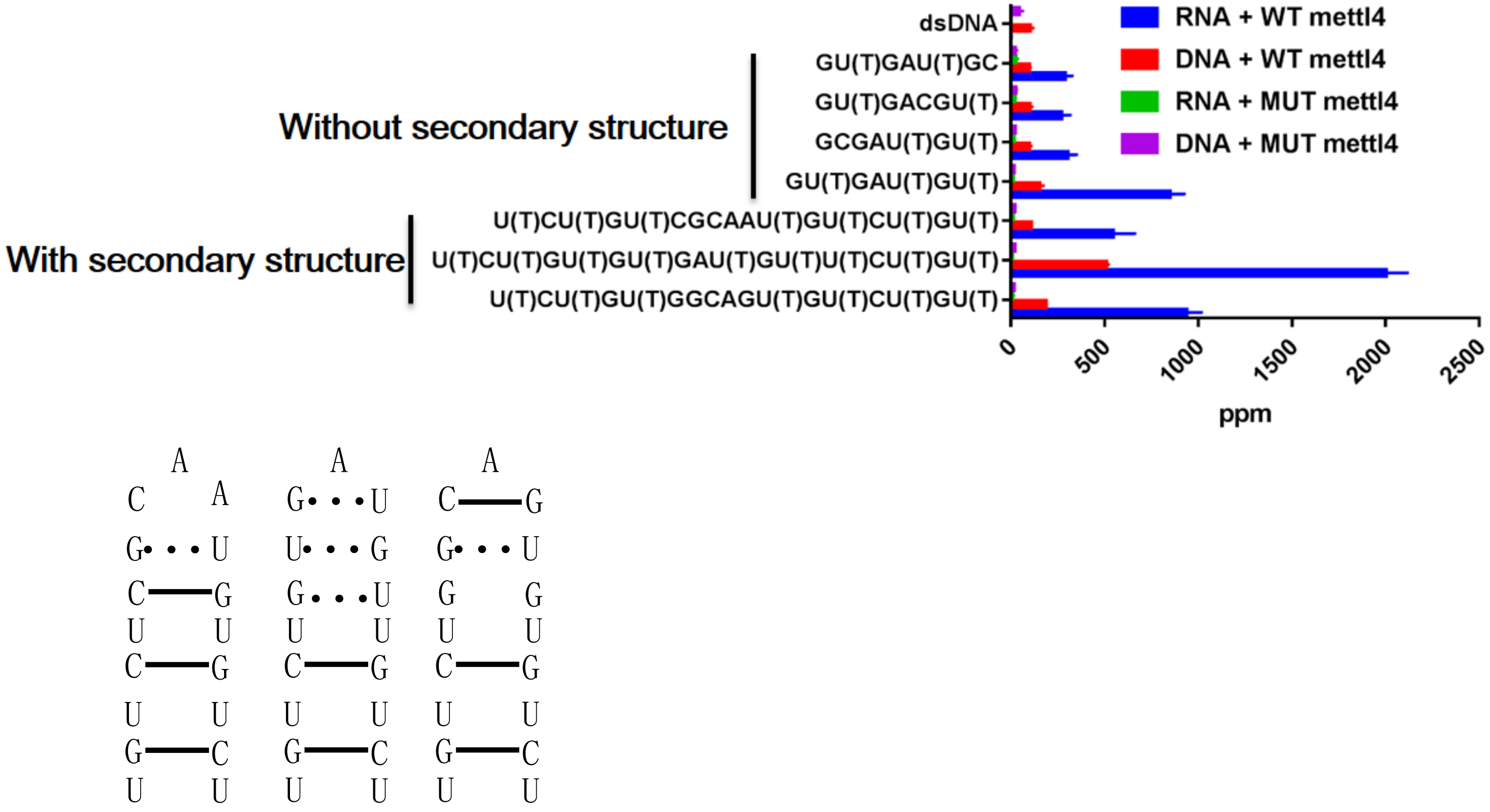

Figure S3 Correlation between biological replicates for the eCLIP-seq samples

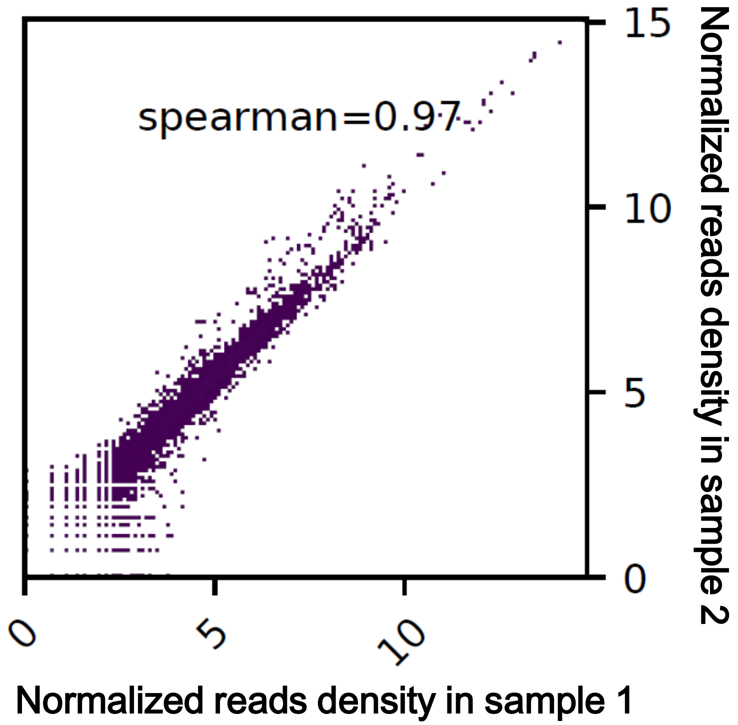

### Figure S4 Generation of knockout Kc cell line

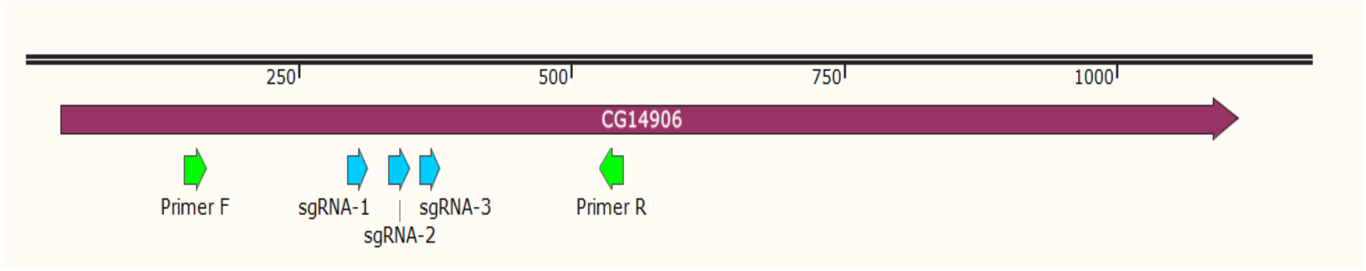


PCR product: 402bp

Small indels produced by single guide RNA caused frame shift mutations.

sgRNA-1: Editing rate: 90%

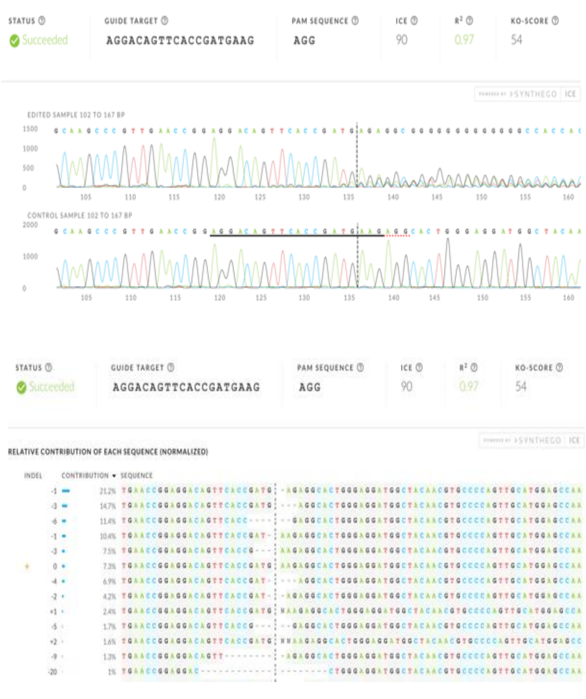


sgRNA-2: Editing rate: 90%

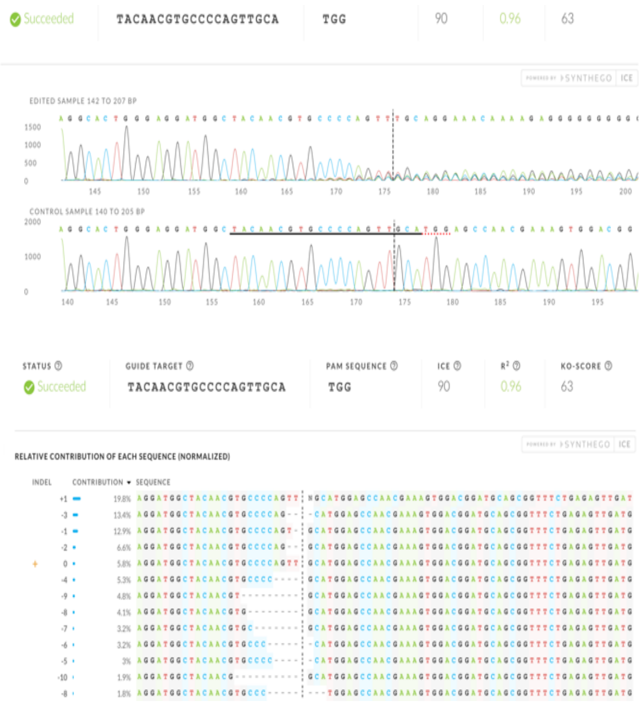


sgRNA-3: Editing rate: 90%

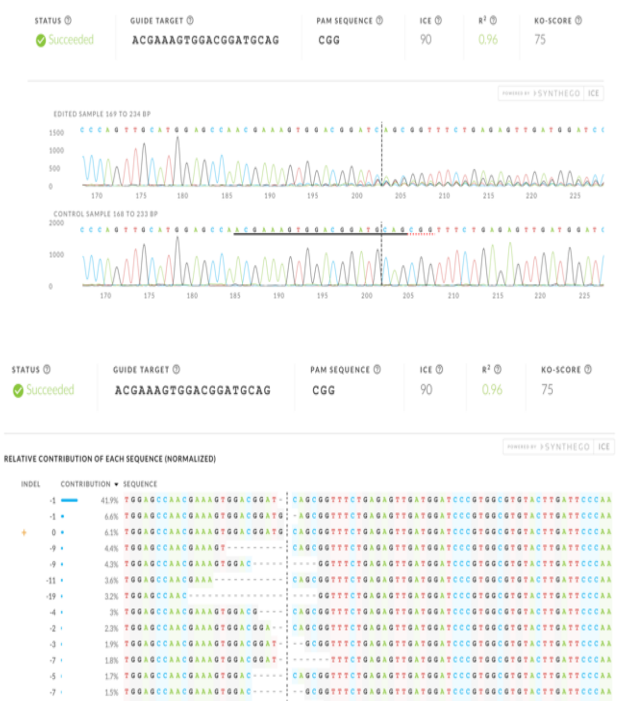

### Figure S5 Rescue of m6A level of U2 snRNA by overexpressing *mettl4* in the *mettl4* KO cells

#### Pentry- PAWF Gateway system:

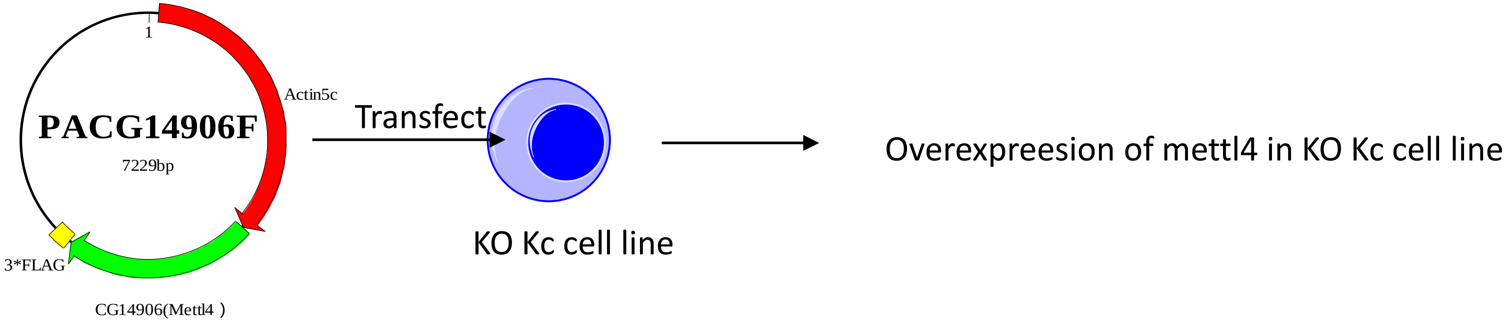


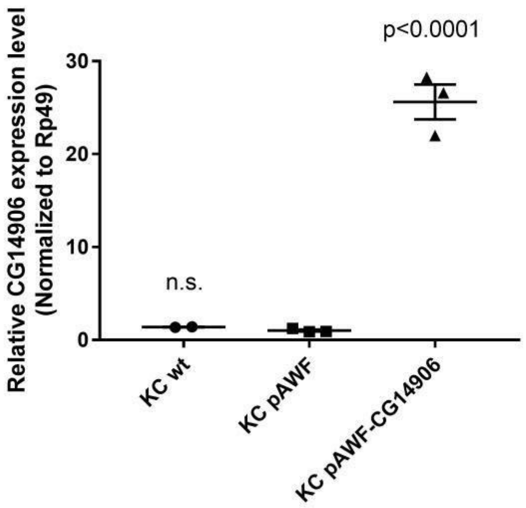


Values are shown as mean±SEM. Results are normalized to KC pAWF.

### Figure S6 Generation of *mettl4* knockout fly

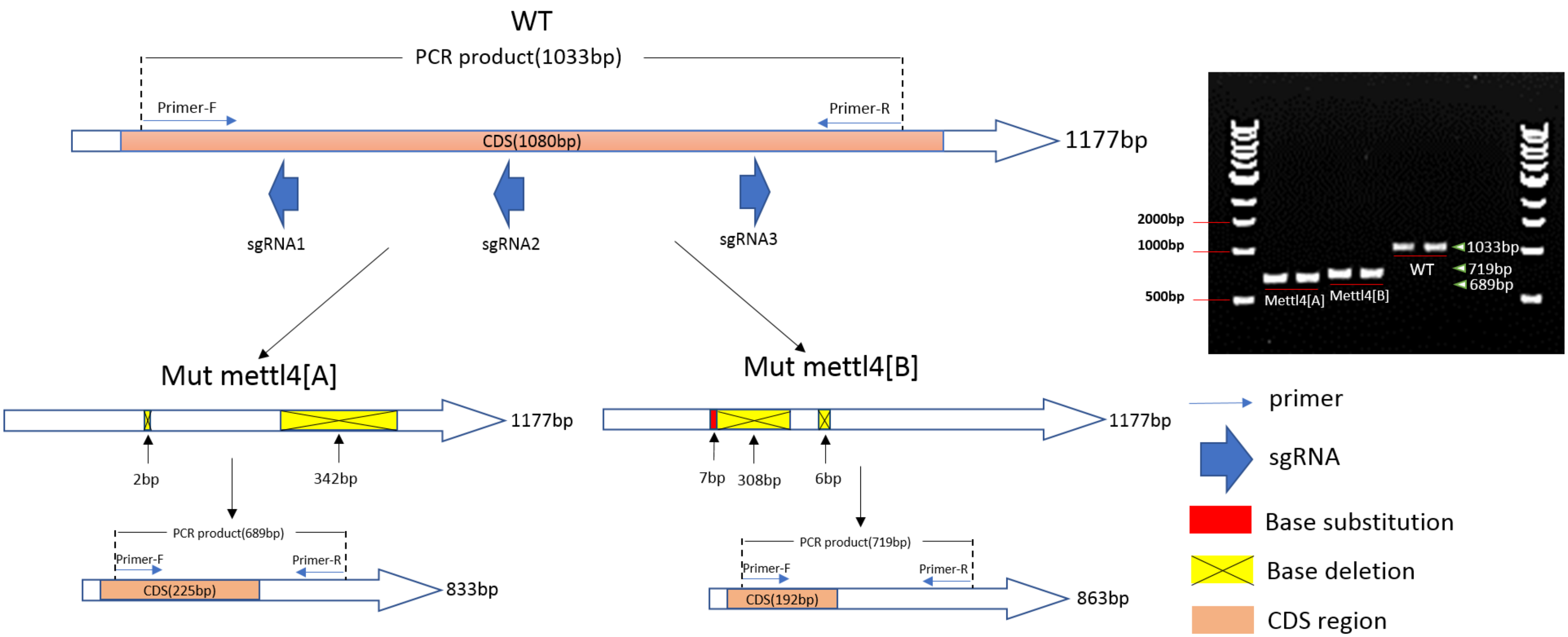

Figure S7 MS spectra of U2 in *mettl4* WT, KO and the rescued cells

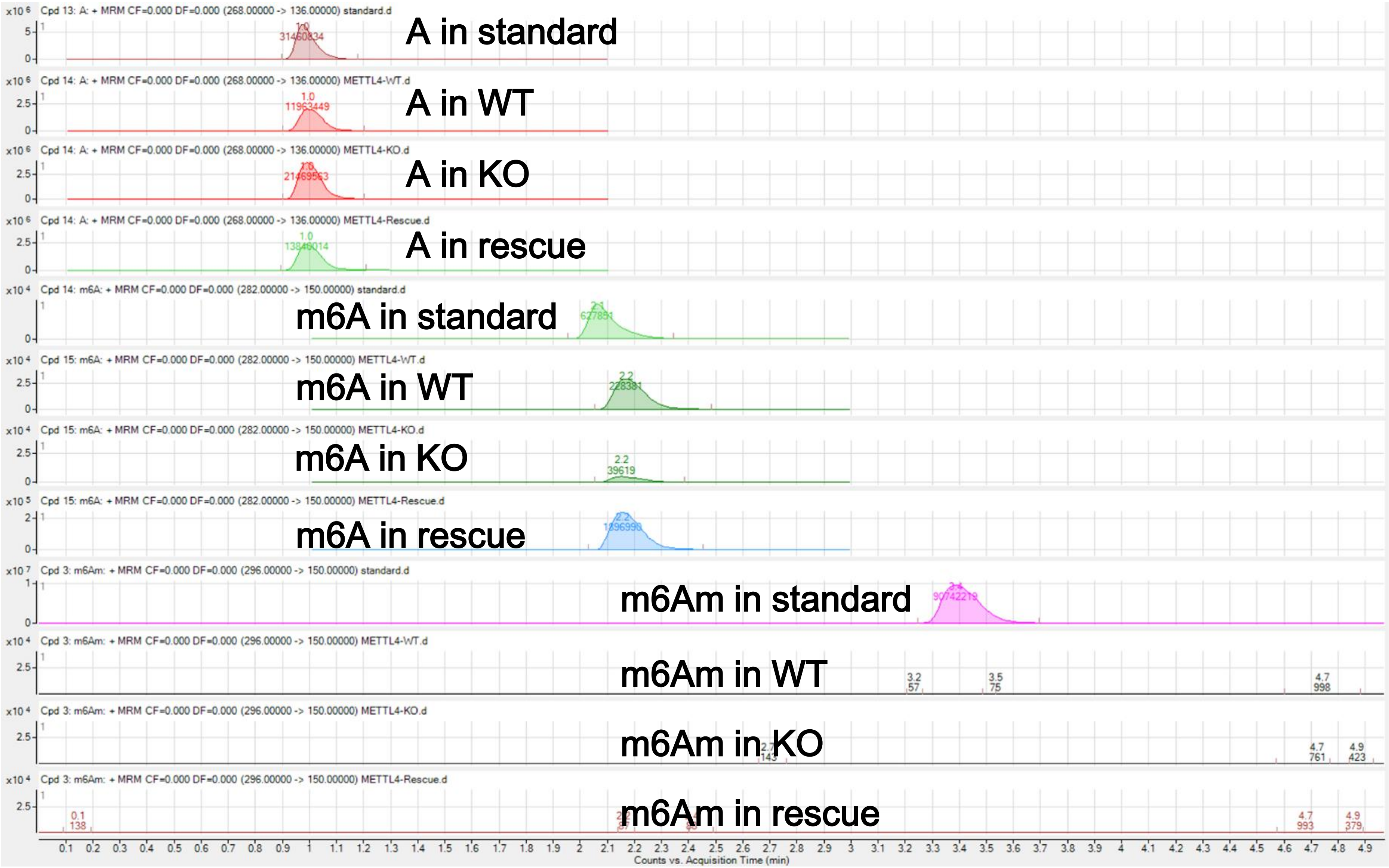

Figure S8 LC-MS/MS results for other independent KO cell lines and flies

Kc cell lines

fly

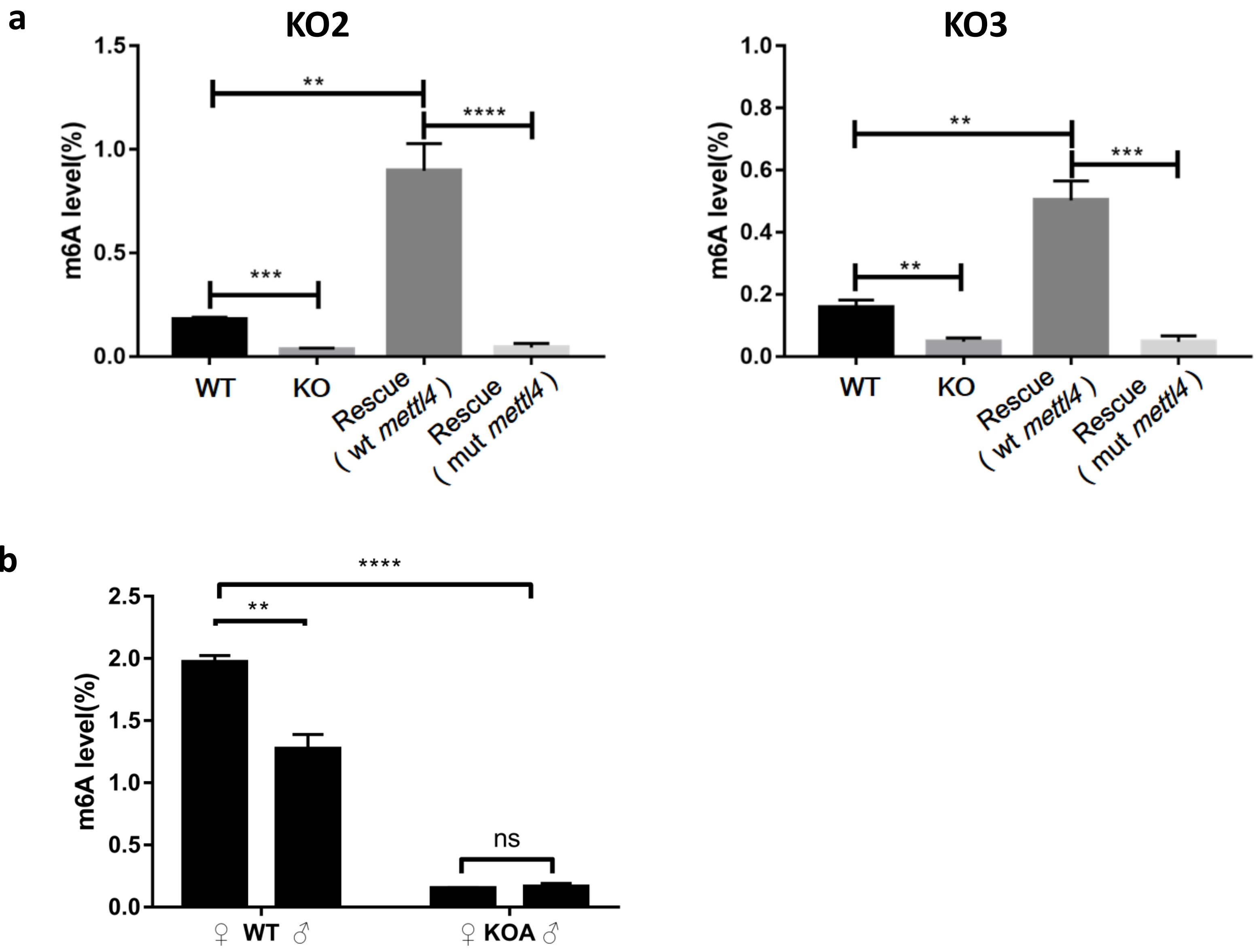

Figure S9 LC-MS/MS results for m6A levels on nuclear and mitochondrial DNA from WT and KO fly cells

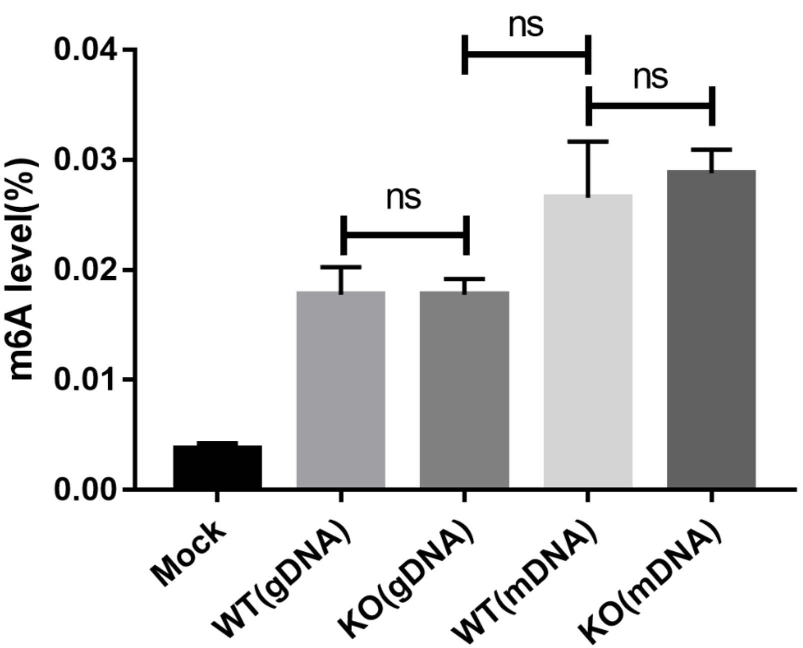
