## Supplementary information for "CG14906 (*mettl4*) mediates m6A methylation of U2 snRNA in *Drosophila*"

Materials and Methods

Protein purification

Cells were harvested and sonicated in lysis buffer (25mM Tris-HCl [pH 8.0], 1M NaCl, 5% glycerol, 2mM dithiothreitol (DTT) and supplemented with protease inhibitors). MBP-tagged METTL4 was purified by affinity for maltose binding protein and the tag was removed by digestion with Tobacco Etch Virus (TEV) protease. Target proteins were further purified by HiTrap Heparin HP columns chromatography, gel filtration chromatography and then was analyzed by SDS-PAGE (Doxtader et al.,2018). The final METTL4 in buffer containing 25 mM Tris (pH 8.0), 150 mM NaCl and 2mM DTT.

Overexpression of Target Proteins

For proteins expressed in E. coli, specified constructs of Drosolhila METTL4 was subcloned into pET28a vector and transformed into Rosetta (DE3) bacterial cells (Novagen). Cells were incubated with agitation at 37℃ until OD600 0.5-0.6, and then cells were induced by adding IPTG to 0.2 mM final concentration. Target proteins were further expressed at 20℃ overnight (Studier,2005).

U2 snRNA purification

Total RNA was extracted from frozen flies with TRIzol reagent according to Invitrogen Life Technologies manual. Streptavidin-conjugated M-280 magnetic Dynabeads (Invitrogen) were used for specific U2 snRNA Isolation. RNase-free beads were washed once with buffer A (10 mM Tris-HCl, pH 7.5, 2 mM EDTA, 2M NaCl), and resuspended in buffer A. Subsequently, biotinylated oligonucleotides were mixed with Dynabeads in buffer A and incubated at room temperature for 30 min with gentle mixing. After the incubation, the oligonucleotide-coated Dynabeads were then washed for four times in buffer B (5 mMTris-HCl, pH 7.5, 1mM EDTA, 1MNaCl) and equilibrated in 6 x SSC solution (1 x SSC is 0.15M NaCl plus 0.015 M sodium citrate, pH 7.0). The oligonucleotide-coated Dynabeads and total RNA in 6 x SSC solutions were heated for 10 min at 75℃. Thereafter, the suspension was incubated at room temperature for 3 hr to allow binding of the U2 snRNAs to the dynabeads. The oligonucleotide-coated Dynabeads were then washed, in succession, three times with 3 x SSC, twice with 1 x SSC, and several times with 0.1 x SSC. U2 snRNA retained on the beads was eluted three times using RNase-free water (Liu et al.,2016)

U2 probe: biotin-atactacactttgatcttagccataaggcc

LC-MS/MS sample preparation and analysis

100ng U2 snRNA was digested with 1U of nuclease P1 (Sigma) in the indicated buffer in 30 uL reaction at 37℃ for 3 hours. Then 1U of Antarctic Phosphatase (NEB) was added to a final reaction volume of 100 μL with phosphatase buffer (NEB) for an hour at 37℃ to dephosphorylate the U2 snRNA. After incubation, 100 μL samples were filtered with Millex-GV 0.22u filters. The samples were run in mobile phase buffer A (water with 0.1% Formic Acid) and 2 to 20% gradient of buffer B (Methanol with 0.1% Formic Acid). MRM transitions were measured for adenosine (268.1 to 136.1, retention time 1.03 min), N6-methyladenosine (m6A) (282.1 to 150.1, retention time 1.79 min), N6,2-O-dimethyladenosine (m6Am) (296.1 to 150.1, retention time 2.40 min). The concentrations of each compound in the samples were calculated using calibration curves constructed with standard compounds of adenosine (Abcam), N6-methyladenosine (Abcam), N6,2'-O-dimethyladenosine (Toronto Research Chemicals). Agilent Mass Hunter LC/MS Data Acquisition Version B.08.00 and Quantitative Analysis Version B.07.01 softwares are used for data collection and analysis.

Kinetics analysis

The kinetic parameters for METTL4 mediated RNA methylation were determined by incubating full-length METTL4 enzyme (300 nM) with increasing concentrations (500 nM – 10 μM) of the substrate (ATCGCTTCTCGGCCTTATGGCTAAGATCAAAGTGTAGTATCTGTTCT) at room temperature in a buffer containing 10 mM HEPES (pH 7.4 @ 25 °C), 5 mM DTT, and 100 μM S-adenosylmethionine (SAM). Aliquots were withdrawn at t = 0, 5, 10, 30, 90 and 150 min and boiled for 3 min to stop all enzymatic activity. The aliquots were further processed according to the LC-MS/MS sample preparation. The levels of the final product (m6A) formed at each time point was determined using mass spectrometry by monitoring the amount of m6A in each aliquot. The observed rate of product formation (kobs) was determined by plotting the concentration of m6A against time for each concentration of the substrate (0.5, 0.75, 1.50, 2.50, 5, 10 μM). The kobs vs substrate concentration curve was fit to the Michaelis-Menten equation for substrate inhibition kinetics using the Graphpad Prism software to obtain the final enzymatic parameters.

eCLIP-seq experiment and data processing

eCLIP-seq experiment was done by Eclipse BioInnovations Company using Sigma M2 anti-FLAG antibody and 10M cells per experiment. Western blot of immunoprecipitation was done during Flag-tag eCLIP in Kc D. Melanogaster cells. 15% of Flag-tag IP, and 1% of input were run on NuPAGE 4-12% Bis-Tris protein gels, transferred to NC membrane, probed with 1:4000 M2 anti-Flag primary antibody (Sigma, #F1804-200UG, lot # SLBT7654) and 1:10000 Mouse TrueBlot ULTRA: Anti-Mouse Ig HRP secondary antibody (Cat # 18-8817-33, Rockland Immunochemical), and imaged with C300 Imager using Azure Radiance ECL. Only the region from 50 to 125 kDa (protein size to

75kDa above) was isolated during eCLIP. Sequencing was performed as SR75 on the HiSeq 4000 platform. Raw sequencing data is processed as descripted previously (Van Nostrand et al.,2016). Briefly, raw fastq data was aligned to the fly reference genome dm6 together with a pool of consensus sequences for rRNA, tRNA, snRNA, snoRNA, miRNA, and LncRNA after trimming adapter and low quality reads. Mapped reads on each region was then converted to reads per million (RPM) for the correction of different sequencing depth among samples. Enrichment score was calculated by $(RPM_{IP}-RPM_{IP\_INPUT})/(RPM_{CONTROL}-RPM_{CONTROL\_INPUT})$.

RNA-seq experiment and data processing

Total RNA was extracted with TRIzol according to the manufacturer's instructions (Invitrogen). PolyA(+) mRNA was isolated from total RNA using the NEBNext® Poly(A) mRNA Magnetic Isolation Module (NEB #E7490). The RNA-seq library preparation was carried out using the NEBNext Ultra II Directional RNA Library Prep Kit (NEB). RNA-seq was carried out on Illumina HiSeq platform with single-end 75bp read length. Raw reads were stripped of adaptor sequences and low quality bases (Q<=20) were removed using Cutadapt (https://cutadapt.readthedocs.io/en/stable/guide.html). The processed reads were aligned to fly genome (dm6) with STAR aligner (version 2.7.0f) and genes with differential alternative splicing between WT and KO cells was identified using RSEM (version 1.3.0). Significantly differentially spliced events are defined as FDR < 0.05 and fold change of exon usage > 2.
